# Embryonic Stem Cell-Specific Responses to DNA Replication Stress

**DOI:** 10.1101/2025.05.16.654332

**Authors:** Ryan C. James, Jerry K. Wang, Siddhanth R. Bhatt, Sophie E. Shadid, Daryl J. Phuong, John C. Schimenti

## Abstract

Genome maintenance is of the utmost importance in stem cells, as mutations can be propagated and cause defects in derivative tissues. Many stem cell types display low mutation rates, with embryonic stem cells (ESCs) being a notable example. The bases for this property are unclear but may be achieved by optimization of various processes including high-fidelity DNA repair, cell cycle checkpoint controls, and hypersensitivity to genotoxic insults that trigger cell death. Here, we investigate the mechanisms underlying the unique responses of mouse ESCs (mESCs) to replication stress (RS) using an array of small molecule inhibitors and genotoxins. We find that whereas mESCs survive under acute RS in an ATR- and CHK1-dependent manner similar to somatic cells, they lack a strong G2/M checkpoint and fail to repair DNA crosslinks in the absence of ATR signaling. Despite the lack of a strong G2/M checkpoint, mESCs maintain a spindle assembly checkpoint (SAC). We posit that mESCs preferentially repair DNA crosslinks in S phase via homology-directed mechanisms, and cells that fail to complete repair before mitosis undergo mitotic catastrophe and cell death. These findings shed light on mutation avoidance mechanisms in ESCs that may extend to other stem cell types.

## Introduction

Mouse ESCs are derived from the inner cell mass of blastocyst stage embryos, which are the progenitors for every cell type in a fully developed mouse^1^. Reflecting their *in vivo* counterparts, ESCs cultured *in vitro* exhibit rapid proliferation, which is driven by constitutive robust expression of factors promoting replication and cell cycle progression^2,3^. Importantly, the pluripotency of ESCs is directly tied to their rapid proliferation, as the activation of checkpoints results in the expression of genes that promote differentiation as well as the downregulation of factors promoting pluripotency^4,5^. This balance of rapid proliferation and retention of pluripotency presents a unique challenge for genome maintenance, and could be expected to result in a high mutation rate^6^. Nevertheless, ESCs have mutation rates that are nearly two orders of magnitude lower than their differentiated counterparts^7^.

ESCs have several properties that likely contribute to their ability to minimize mutation accumulation. One important mechanism is the use of high-fidelity DNA repair pathways, such as homology-directed repair (HDR). ESCs exhibit high expression of homologous recombination (HR) factors and spend most of the cell cycle in S phase, during which HDR is preferentially employed ^2,8,9^. Importantly, ESCs lack a G1/S checkpoint due to attenuated p53 signaling^10^, a feature associated with hypersensitivity to genotoxic insults such as ionizing radiation^11^. Counterintuitively, the lack of this checkpoint has been proposed as a mutation avoidance strategy: by allowing cells with DNA damage to undergo catastrophe rather than attempting repair upon entrance into S phase, genetically compromised cells can be removed from the pool^12^.

While the responses of ESCs to induced strand breaks highlight mutation avoidance strategies, replication errors are believed to be the predominant source of spontaneous mutations. Mouse ESCs experience constitutive RS that is attributed to short gap phases^6^. Understanding how ESCs respond to RS is therefore critical for elucidating mechanisms of mutation avoidance. Most studies on responses of ESCs to RS have used genotoxic chemotherapeutics such as cisplatin and Adriamycin (doxorubicin)^10,13,14^. Cisplatin induces intra- and interstrand crosslinks that impede CMG helicase progression^15^, while doxorubicin intercalates into DNA and inhibits topoisomerase II, resulting in strand breaks and replication fork stalling^15^. In response to DNA crosslinks induced by agents such as mitomycin C (MMC), mouse ESCs display phosphorylation of proteins at sites modified by ATR and/or ATM, the apical DNA damage response kinases^13^. However, cisplatin treatment elicits neither the phosphorylation nor activity of CHK1 (checkpoint kinase 1, formally CHEK1), a key downstream effector of ATR^13,14^. By contrast, in somatic and cancer cells, cisplatin-induced CHK1 activation mediates the G1/S, intra-S, and G2/M checkpoints via downstream inhibitory phosphorylation of CDK1 (cyclin dependent kinase 1)^13,14,16,17^. Although CHK1 is not activated in ESCs following cisplatin or doxorubicin treatment, these agents do induce G2 accumulation prior to extensive cell death^10,14^. This suggests that ESCs may possess an intact spindle assembly checkpoint (SAC), though direct evidence is currently lacking.

Here, we investigate the mechanisms underlying the unique responses of ESCs to RS using an array of small molecule inhibitors and genotoxins. We focused on conditions that trigger checkpoint activation in ESCs, as well as the downstream signaling and transcriptional changes induced by DNA crosslinking agents. These studies identified a critical role for the SAC in compensating for the absence of the G2/M checkpoint, as well as cell type-specific functions of ATM and ATR in managing crosslink-induced stress.

## Results

### Responses of mouse ESCs (mESCs) to replication stress

To probe checkpoint activation in mESCs, we selected a panel of genotoxins that induce RS via distinct mechanisms. Bleomycin, an ionizing radiation (IR) mimetic, was used to induce strand breaks^18^. MMC was used to induce inter- and intrastrand crosslinks^19^, as was Melphalan, an agent that induces a higher proportion of interstrand crosslinks^20^. Finally, aphidicolin, a selective inhibitor of replicative DNA polymerases, was utilized at a dose (1µM) sufficient to stall DNA replication.

Treatment of mESCs with MMC or melphalan induced little activation of CHK1 (phosphorylation of S345) or inhibition of cyclin dependent kinase 1 (CDK1) (phosphorylation of Thr14, Tyr15; Figure 1A), but MMC nevertheless induced G2/M accumulation (Figure 1C). In contrast, MMC-treated MEFs (mouse embryonic fibroblasts) exhibited robust CHK1 phosphorylation (Figure 1B) but only modest G2/M accumulation (Figure 1D). These responses are consistent with previous observations in human and mouse cells treated with various crosslinking agents, including lack of CHK1 activation in cisplatin-treated human ESCs (hESCs)^19^,^21,22^. Aphidicolin treatment induced phosphorylation of CHK1 and CDK1 in both mESCs and MEFs (Figure 1A,B), causing cultures of both cell types to accumulate in S phase (Figure 1C,D) and undergo a ∼50% reduction in viable cells after 24-hours (Figure S1A). These results suggest that MEFs and ESCs respond differently to crosslinks, yet similarly to acute RS.

**Figure 1.**
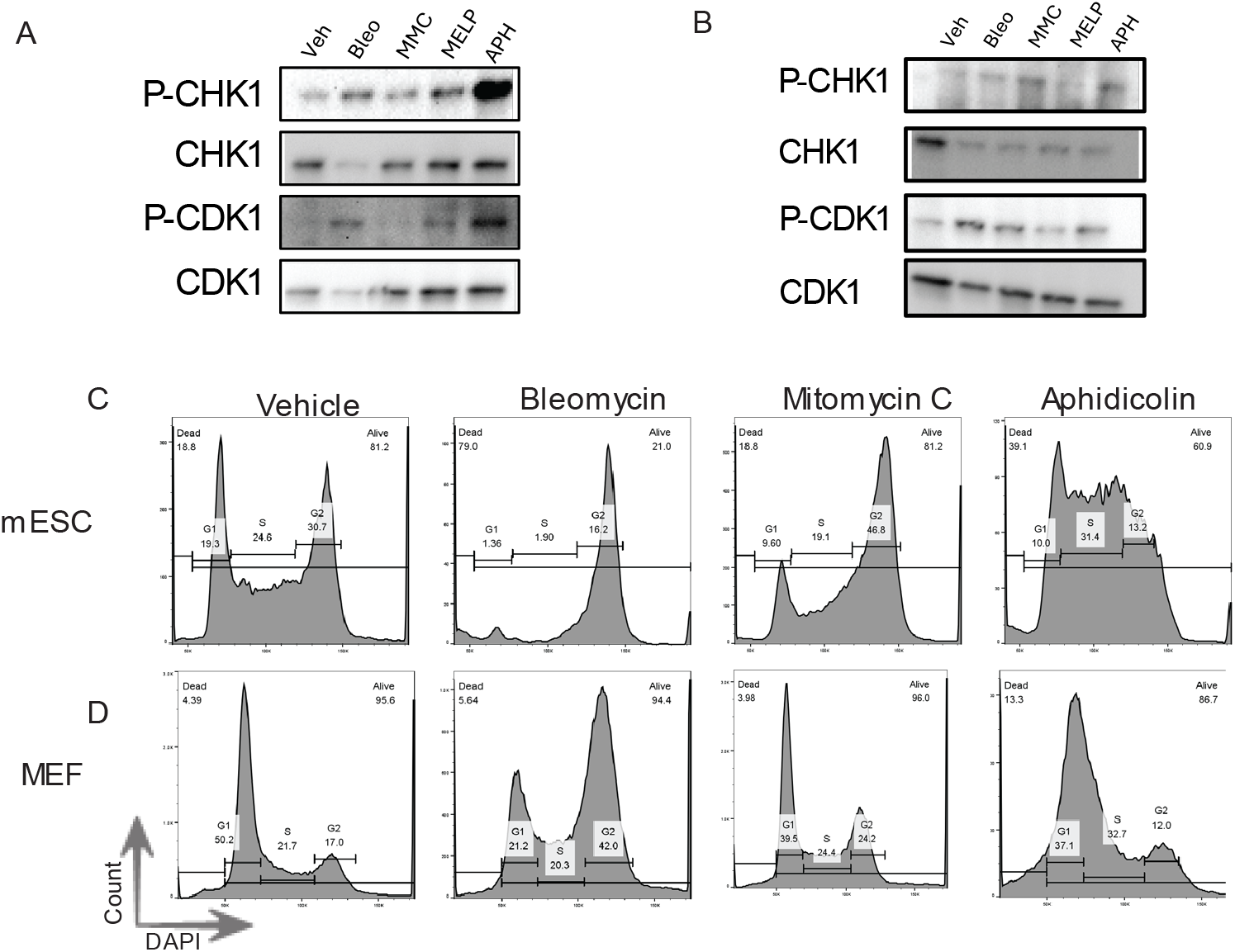
Mouse ESC and MEF responses to agents that induce replication stress. (A) Western blots for p-CHK1 (S345), CHK1, p-CDK1 (Thr14, Tyr15) of mESCs or MEFs (B) treated with indicated drugs for 24 hours. (C) DNA content analysis of mESCs or MEFs (D) treated with the indicated drugs for 24 hours. Veh, vehicle; Bleo, bleomycin; MMC, mitomycin C; MELP, melphalan; APH, aphidicolin.

### Mouse ESCs retain an ATR- and CHK1-mediated intra-S phase checkpoint

Given that aphidicolin-induced RS triggered robust CHK1 phosphorylation and S-phase accumulation in mESCs, we tested whether CHK1 had a functional role in mediating the accumulation and survival of these cells in S-phase. Indeed, co-treatment with the CHK1 inhibitor CHIR-124 (CHK1i) reduced APH-induced S phase accumulation by ∼40%(Figure 2A, Figure S3A). This effect was also observed in MEFs (Figure 2B, Figure S3B). In both mESCs and MEFs, co-treatment resulted in a dramatic increase in the sub-G1 population, approximately accounting for the lost S-phase population. The marked cell death as a result of this co-treatment was confirmed with viability assays.

**Figure 2.**
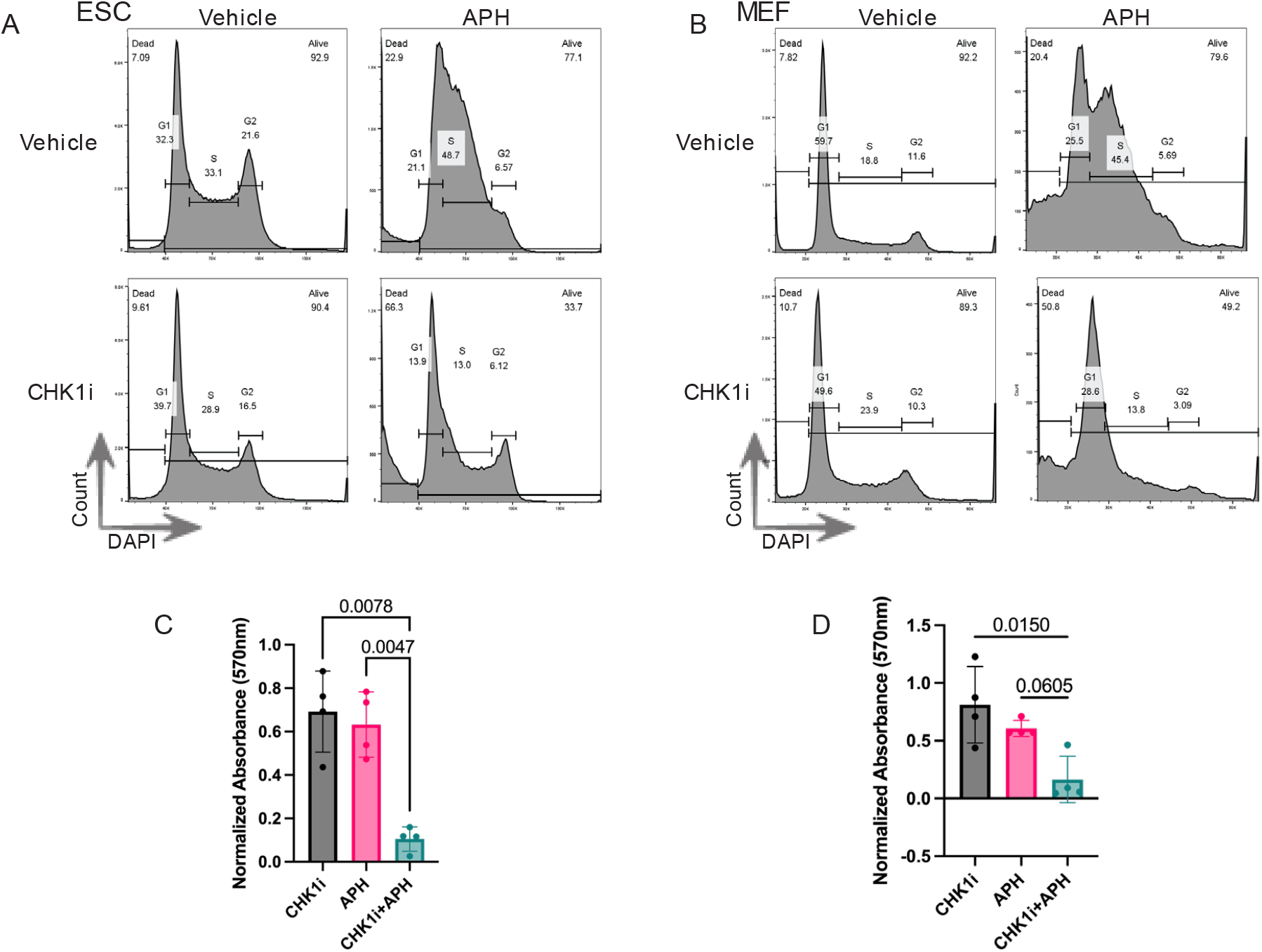
Survival of ESCs under acute replication stress is CHK1-dependent. (A) DNA content analysis of mESCs or (B) MEFs treated with the indicated drugs for 24 hours. Drugs were added simultaneously. (C) Viability analysis via MTT assay of mESCs or (D) MEFs. MTT intensity is normalized to an untreated sample. Error bars show SD.

This analysis demonstrated a >90% reduction in the surviving populations of mESCs and MEFs after 24 hours of co-treatment with CHK1i and APH (Figure 2C,D).

The similar responses of mESCs and MEFs to RS led us to hypothesize that ESCs mount a canonical RS response, specifically ATR-mediated phosphorylation (activation) of CHK1^23^. Supporting this, inhibition of ATR with VE-822^24^ produced effects comparable to CHK1 inhibition in both mESCs and MEFs co-treated with aphidicolin (Figure S2A,B). Additionally, co-treatment of mESCs with a DNA-PKcs inhibitor - previously shown to enhance intra-S phase checkpoint activation^25^ - led to a marked reduction in ESC viability (Figure S2E), further indicating that ESCs rely on canonical checkpoint pathways to cope with RS.

### The role of the G2/M checkpoint in responding to crosslinks in mESCs

To determine whether MMC treatment elicits a DNA damage response (DDR) in mESCs despite the absence of activated CHK1, we immunostained MMC-treated mESCs for γH2A.X, a marker of DNA damage and RS that is phosphorylated by the apical DDR kinases, ATR and ATM^26^ The MMC-treated cells exhibited a dramatic and significant increase in γH2A.X nuclear foci and intensity compared to untreated cells (Figure 3B), indicating active upstream DDR signaling.

**Figure 3.**
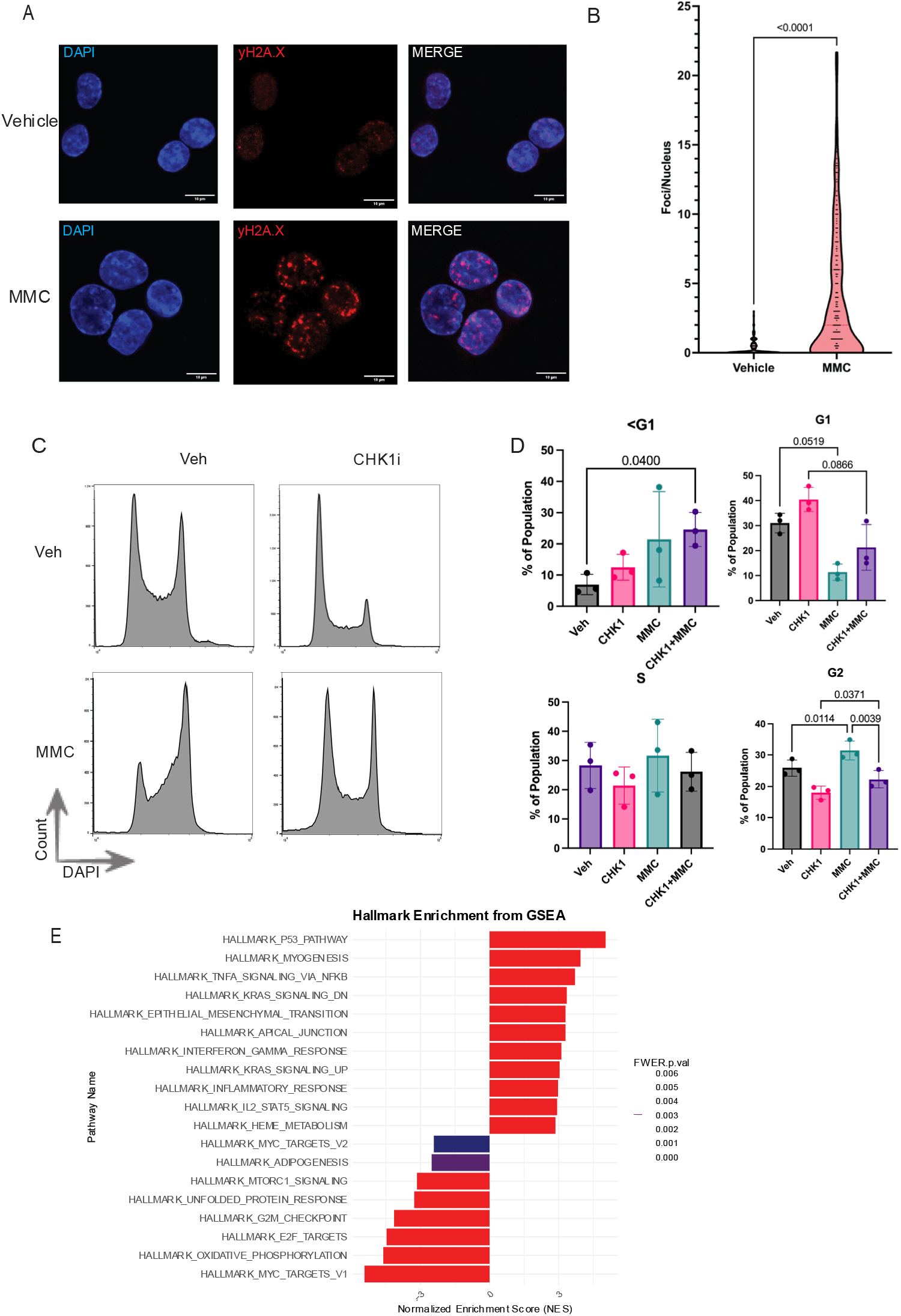
The response of mESCs to crosslinks Is not CHK1-dependent. (A) Representative images of γH2A.X staining of mESCs treated with MMC for 24 hours. (B) Quantification of γH2A.X foci per nuclear slice. Z=3, n>150, N=3. (C) DNA content analysis of mESCs treated with the indicated drugs for 24 hours. Drugs were added simultaneously. (D) Quantification of C. Error bars show SD. (E) Hallmark enrichment plot (MMC-treated vs Vehicle) of the top and bottom 10 hallmarks from RNA sequencing of mESCs treated with MMC for 24 hours.

Given that CHK1 was not phosphorylated—and therefore not activated—in response to DNA crosslinking agents, we hypothesized that CHK1 inhibition would have minimal impact on the response of mouse ESCs to MMC. Consistently, CHIR-124 treatment alone caused an increase in Sub-G1 and G1 phase populations and a decrease in S and G2 populations (Figure 3C). In contrast, MMC treatment alone induced G2 accumulation, as expected (Figure 3C). Remarkably, co-treatment with CHIR-124 and MMC produced additive effects on cell cycle distribution (Figure 3C,D). To assess whether CHK1 inhibition altered the cellular response to MMC, we normalized the co-treated samples to their respective single-agent controls (e.g., CHK1i+MMC normalized to MMC alone). This analysis revealed no significant effect of CHK1 inhibition on MMC-induced cell cycle changes (Figure S6). Similarly, CHK1 inhibition did not alter MMC-induced cytotoxicity, as determined by survival assays (Figure S4D). Together, these findings suggest that the reported roles of CHK1 in replication origin firing, fork stabilization, and checkpoint activation do not substantially influence the response of mESCs to DNA crosslinks^27^.

The apparent lack of CHK1-mediated checkpoint activity in MMC-treated mESCs led us to hypothesize that transcriptional changes associated with checkpoint activation would also be absent. To test this, we performed RNA sequencing on MMC-treated mESCs and conducted gene set enrichment analysis (GSEA) of hallmark pathways^28^. As expected, the reduction in S-phase entry correlated with decreases in the “MYC Targets” and “E2F Targets” hallmarks. Interestingly, there were indications of MMC-induced differentiation. For example, we observed >2-fold increases in the expression of EGFR, FGFR2, and FGFR4 (Figure S5E)^29,30^, downregulation of the key pluripotency regulators OCT4 and NANOG (Figure S5F)^31,32^, and differential expression of many aldehyde dehydrogenase genes, which is observed in the differentiation of many stem cell types (Figure S5D)^33^.

Most striking was an unexpected decrease in the “G2M Checkpoint” hallmark, reflecting fewer cells progressing through this phase of the cell cycle (Figure 3E). As WEE1 acts downstream of CHK1 and phosphorylates CDK1 to activate the G2/M checkpoint, we expected its inhibition to also have little effect on mESCs treated with MMC^34^. Indeed, we inhibited WEE1 with MK-1775 (WEE1i) and observed no change in DNA content profiles between MMC-treated and co-treated samples (Figure S4A)^35^. Unlike treatment with CHK1i, inhibition of WEE1 did not reduce S or G2 populations (Figure S4A). Further, treatment with WEE1i also did not sensitize mESCs to MMC as observed via survival assay (Figure S4C).

To probe the role of a G2/M checkpoint in MEFs, we inhibited WEE1 in combination with MMC treatment. Notably, DNA content analysis showed that WEE1i elicited a decrease in the G1 population and an increase in the G2 population (Figure S4B). This treatment also induced a decrease in p-CDK1 levels (Figure S4E). Further, survival analysis showed that co-treatment of MEFs with MMC plus either CHK1i or WEE1i resulted in a modest reduction in viability, similar to that seen in mESCs treated with MMC alone (Figure S4C). In sum, unlike somatic cells such as MEFs exposed to crosslink-induced RS, ESCs appear to lack a robust G2/M checkpoint.

### Spindle Assembly Checkpoint signaling mediates cell cycle arrest in response to crosslinks in mESCs

Given the paradoxical observation of G2 accumulation in the absence of classical G2/M checkpoint signaling, we investigated whether the spindle assembly checkpoint (SAC) might mediate this arrest. To test this, we inhibited the key SAC regulator MPS1^36,37^ using a small molecule inhibitor MPI-0479605 (MPS1i). MPS1i treatment alone had minimal impact on mESC cell cycle progression. However, co-treatment with MPS1i and MMC abolished the G2 accumulation observed in MMC-treated mESCs and led to a substantial increase in the sub-G1 population (Figure 4A,B). This was accompanied by a 40% decrease in cell viability (Figure 4C). In contrast, MPS1 inhibition in MEFs, with or without MMC, had no significant effect on cell cycle distribution (Figure S7A).

**Figure 4.**
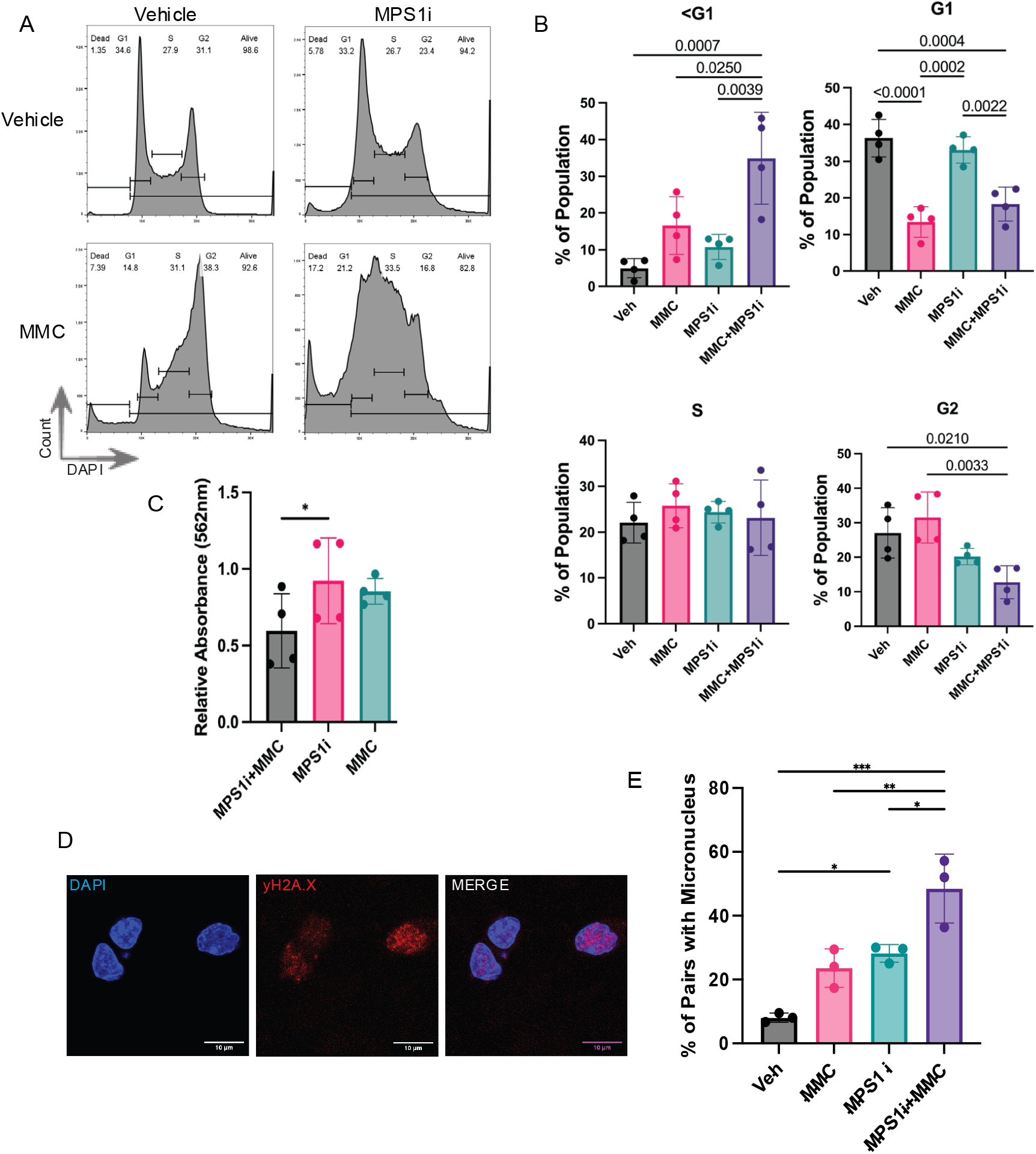
ESCs rely on the SAC to prevent chromosomal abnormalities in response to replication stress. (A) DNA content analysis of mESCs treated with the indicated drugs for 24 hours. Drugs were added simultaneously. (B) Quantification of A. Error Bars show SD. (C) Viability analysis via MTT assay of mESCs treated with the indicated drugs. (D) Representative image of micronucleus formed during treatment with MPS1i+MMC. (E) Quantification of D shows the %of pairs of cells with a micronucleus between them. 3 biological replicates with 20 pairs analyzed each.

As inhibition of the SAC ablated the MMC-induced accumulation of mESCs in G2, we asked whether the remaining cells would display mitotic abnormalities in the form of elevated micronuclei (Figure 4D). Remarkably, ∼half of MMC- and MPS1i-treated cell pairs displayed micronuclei, a significant increase compared to MPS1i or MMC alone (Figure 4E), as well as a significant increase in γH2A.X foci over MMC alone (Figure S8). These findings demonstrate that ESCs have an intact SAC that plays a role in reducing mitotic abnormalities and DNA damage caused by blocks to DNA replication. This SAC is mediated by canonical MPS1 signaling and is responsible for the transient accumulation of mESCs in G2 in response to crosslinks. This reinforces our findings that the G2/M checkpoint is not responsible for the G2 accumulation of mESCs treated with crosslinking agents.

### Differential regulation of checkpoints by apical DNA damage response kinases in MEFs and ESCs

Considering that mESCs spend the majority of their cell cycle in S phase and preferentially utilize HDR compared to differentiated cells, we suspected that the regulation of checkpoint responses in mESCs would be more ATR-than ATM-dependent. Since ATR is a crucial mediator of many crosslink repair pathways, its inhibition is used as an adjuvant to DNA crosslinking drug treatments^38–40^. To investigate the role of ATR in the response of mESCs to DNA crosslinks, we treated mESCs with VE-822 (ATRi) or KU-60019 (ATMi) in conjunction with MMC. Expectedly, co-treatment of MEFs and mESCs with MMC and ATRi resulted in a viability decrease compared to ATRi alone (Figure 5A,B). Importantly, MEFs were sensitized to MMC when co-treated with ATMi, while this effect was not observed in mESCs (Figure 5A,B). Additionally, treatment with ATRi and ATMi failed to induce further cell death in mESCs compared to ATRi alone.

**Figure 5.**
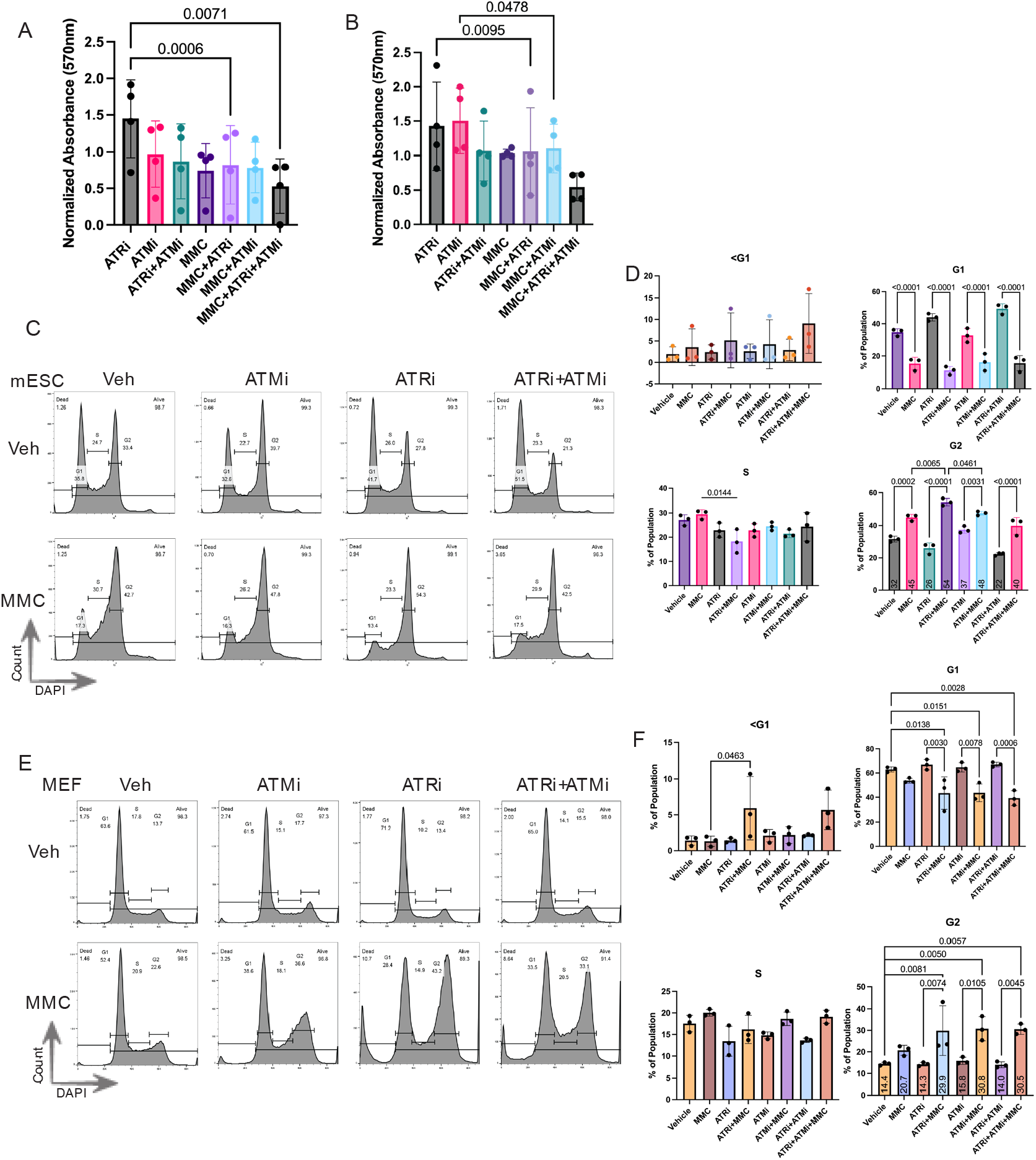
Differential responses of ESCs and MEFs to the inhibition of ATR and ATM. (A) Viability analysis via MTT of mESCs or MEFs (B) treated with the indicated drugs. (C) Flow cytometry analysis of mESCs treated with indicated drugs. (D) Quantification of C. (E) Flow cytometry analysis of MEFs treated with indicated drugs. (F) Quantification of C. All error bars show SD

DNA content analysis of both cell types treated with these combinations was also revealing. In MEFs without insult, treatment with ATRi, ATMi or both had no discernible effect (Figure 5E). ESCs treated with MMC+ATRi accumulated in G2 to an even greater extent than MMC alone (Figure 5C,D), but MMC+ATMi did not result in any change beyond those induced by MMC (Figure 5C,D). Concordant with survival assay data, co-treatment of mESCs with both ATR and ATM inhibitors did not result in a significant change in any population compared to co-treatment with just ATRi (Figure 5D). In contrast, MMC co-treatment of MEFs with ATRi, ATMi or both induced a decrease in G1 population and an increase in G2 (Figure 5E,F).

Finally, to probe the cell type-specific reliance on these kinases for the activation of checkpoints, we analyzed CDK1 phosphorylation in MEFs and mESCs treated with ATR and ATM inhibitors in addition to MMC. As expected, we observed that ATR inhibition in the presence or absence of MMC greatly reduced p-CDK1 levels in both cell types; however, ATM inhibition could ablate p-CDK1 levels in MEFs but not mESCs (Figure S9C,D). Taken together, these results indicated that ATM is somewhat dispensable for ESCs in response to crosslinks, though it plays a role in differentiated cell types. Further, the inhibition of ATR downregulated the G2/M checkpoint, yet enhanced MMC-induced G2 accumulation in both cell types. This reinforces our model that the G2/M checkpoint is not responsible for the G2 accumulation of mESCs in response to crosslinks.

## Discussion

Cells at any developmental stage must balance genomic integrity with a cell type-specific array of challenges. In embryonic stem cells, these challenges include rapid proliferation, constitutive replication stress (RS), and epigenetic states specific to pluripotency^3,6,10,41^. Through a study of checkpoint and repair responses, we demonstrate several foundational aspects of genome maintenance in murine embryonic stem cells (mESCs).

Given the lack of a G1/S checkpoint in mESCs, and the lack of checkpoint responses to other insults to replication such as crosslinks, we did not expect to observe an intra-S phase checkpoint response. For example, a study of human ESCs did not observe such a response to RS induced by thymidine replacement^14^. However, we found that mESCs do respond in a CHK1-dependent manner to replication stress caused by Aphidicolin treatment. Given the rapid proliferation of ESCs, these cells are likely to undergo transient nucleotide depletion, necessitating responses that promote fork stabilization or a cessation of replication before DNA synthesis is resumed. This is supported by the knowledge that both ATR and CHK1 are essential genes, with the loss of either resulting in early embryonic lethality^23,42^.

Despite having the capacity to activate ATR, CHK1, and downstream checkpoints in response to ssDNA accumulation, we have demonstrated that mESCs, like hESCs, do not activate CHK1-dependent signaling in response to crosslinks^14^. By blocking G2/M checkpoint activation through ATR, CHK1, or WEE1 inhibition, we reveal that this checkpoint is dispensable in mESCs, in contrast to differentiated cells. Despite the inhibition of G2/M checkpoint mediators, we observed that ESCs treated with MMC accumulated in G2.This has been observed in response to similar agents, such as cisplatin and doxorubicin^10,13^. Importantly, we find that this accumulation can be abrogated by inhibiting MPS1, a key mediator of the SAC^37,43^. In this condition, cells that survive are prone to mitotic abnormalities, highlighting the reliance on a functional SAC in the absence of a G2/M checkpoint.

Previous findings indicate that CHK1 localizes to centrosomes in mESCs, which would result in limited activation of checkpoints in response to certain types of stress^44^. Adding to this, our findings support the existence of a threshold at which ATR begins to phosphorylate sequestered CHK1. The limited ssDNA exposure induced by crosslinks and crosslink repair may result in local ATR activation, allowing mediation of repair without affecting CHK1. At the other extreme, the ssDNA exposure induced by aphidicolin would result in widespread ATR activation, at which point the sequestration of CHK1 is insufficient to prevent its activation.

Beyond the sequestration of CHK1, the inability of mESCs to activate the G1/S or G2/M checkpoint in response to genotoxic insult is mechanistically rooted in the constitutively high expression of factors that promote cell cycle progression as well as limited expression of CDK inhibitors such as p21^45–47^. As such, the ability of mESCs to reach a threshold of CDK inhibition necessary for checkpoint activation is limited by the sheer quantity of CDK expression and the overexpression of CDC25 phosphatases that antagonize checkpoint activation^48,49^.

Our finding that ATM plays a differential role in response to crosslinks in MEFs and mESCs indicates that the length of the G1 phase impacts the response to crosslinks. Crosslinks can distort the DNA double helix, allowing recognition and processing by mismatch repair proteins, including MLH1^50^. As MLH1 interacts with ATM, this recognition and processing results in checkpoint activation before S phase^51^. In mESCs, a short G1 phase and an inability to activate a G1/S checkpoint reduces the time in which crosslinks can be recognized and processed by MMR factors. As such, mESCs must react to replication forks that encounter unprocessed ICLs, necessitating ATR-mediated responses. Importantly, though we observe phosphorylation of H2A.X in response to MMC, our data could also suggest that crosslinks go entirely unrepaired in ESCs. The cotreatment of ESCs with MMC and ATRi increased G2 accumulation, rather than greatly increasing cell death, which could be the result of an accelerated cell cycle and an increase in cells arrested at the SAC.

Regarding the function of this cell type, the suppression of checkpoints promotes rapid proliferation as well as the maintenance of pluripotency, as the inhibition of cell cycle progression results in differentiation^4,52^. Failing to activate a G1/S checkpoint is proposed as a mutation avoidance strategy that preserves rapid proliferation while culling cells with excess damage^11^. With our findings, it is tempting to speculate that the lack of a G2/M checkpoint follows the same model. Of note, we observe that ESCs treated with MMC display transcriptional changes consistent with differentiation. This is consistent with the tendency of other stem cell types to differentiate in response to DNA damage to prevent the persistence of mutations in a self-renewing population^53,54^.

Taken together, our findings provide foundational knowledge in the understanding of the cell cycle in mammalian pluripotent stem cells. We show that the response to RS in mESCs is unique from that of multipotent cells and is differentially activated by certain DNA damaging agents. These findings provide key insights into the potential mutation avoidance strategies of this cell type, some of which may be mirrored in other stem cell lineages.

## Methods

### Cell culture and drug treatments

B6(Cg)-Tyr(c-2J)/J-PRX mouse embryonic stem cells (mESCs) from The Jackson Laboratory were cultured in 2i+LIF media in 6-well plates coated with Laminin^55^. Cells in 2i+LIF were passaged 1:8 every 4 days and were not cultured beyond 20 passages. Media changes were performed daily. SV40 Large T antigen-immortalized Mouse Embryonic Fibroblasts (MEFS) were cultured in medium containing 10% FBS, 1% Penicillin-Streptomycin, and 1X NEAA (Gibco). MEFs were passaged 1:6 every 4 days.

Unless otherwise noted, treatment of cells was performed for 24 hours with drugs diluted in culture media. Concentrations of genotoxins and inhibitors used can be found in Table 1. mESCs were treated 2 days after passage and MEFs were treated 3 days after passage, and treatments were performed in 6-well plates.

**Table 1.**
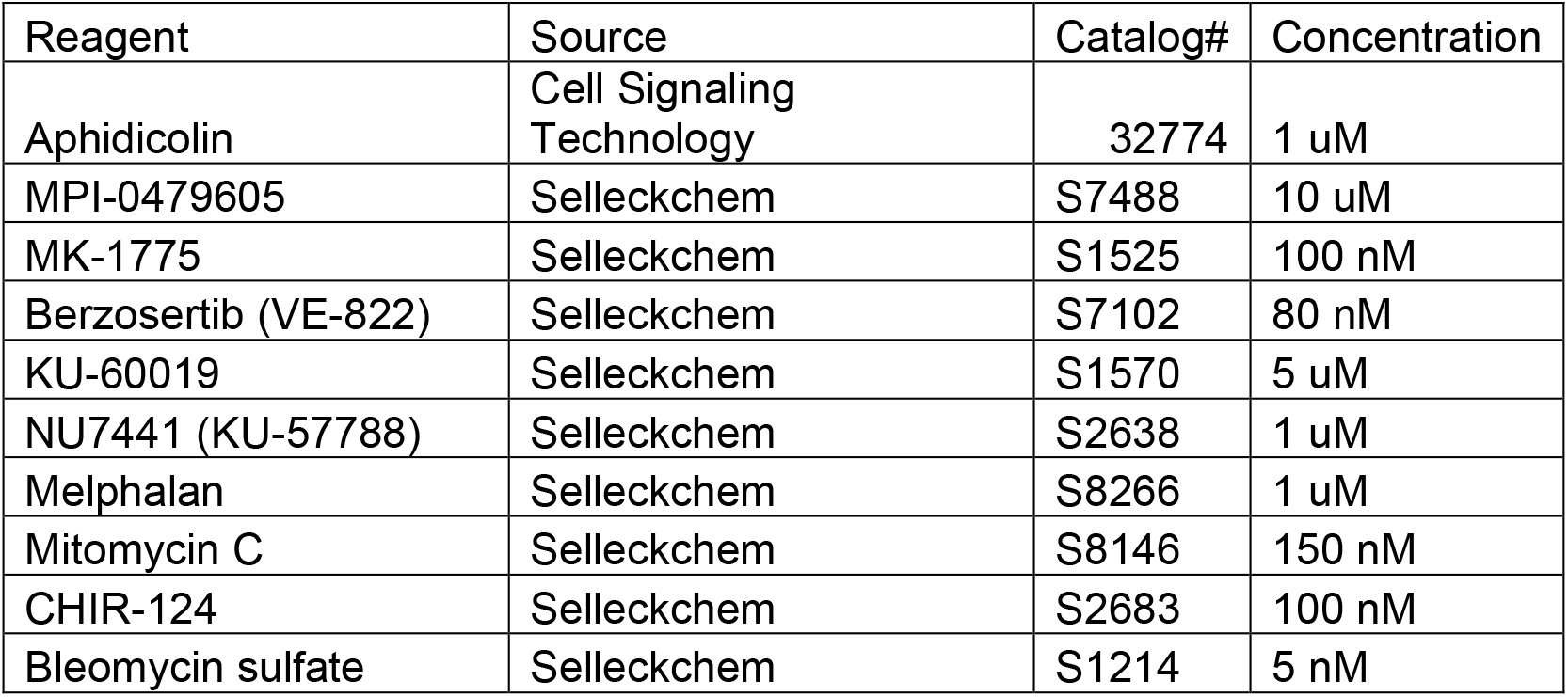
Genotoxins and kinase inhibitors employed in the study. Final concentrations used to treat cells are listed.

### DNA Content Analysis via Flow Cytometry

After treatment, all cell types were trypsinized, suspended in serum-containing media, and pelleted. Following a wash with 1X PBS, cells were fixed with 70% ethanol for a minimum of 24 hours. Subsequently, cells were pelleted, washed with PBS containing 1% BSA (Flow buffer), pelleted once again, and resuspended in Flow buffer containing 10 mg/ml RNAse A and 1 µg/mL DAPI. Cells were strained with a 40 µm filter before flow cytometry on a ThermoFisher Attune NxT Flow Cytometer. Analysis of populations was performed using FlowJo software.

### Western blotting

Cell pellets were resuspended in RIPA then frozen to assist lysis. Protein concentrations were determined using BCA assay (ThermoFisher). Denatured lysates were separated in 4-20% polyacrylamide gels and wet transfer was performed at 1000mA for 1 hour using Bolt-Mahoney transfer buffer. Membranes were blocked in 5% BSA at RT for 1 hour before overnight incubation at 4C with primary antibodies: Rabbit Phospho-CDK1 (Thr14, Tyr15) polyclonal (Invitrogen 44-686G), Rabbit Phospho-CHK1 (Ser345) (133D3) (Cell Signaling Technology 2348), Mouse CHK1 monoclonal (Invitrogen MA1-91087), Mouse CDK1 Monoclonal Antibody (A17) (Invitrogen 33-1800), Mouse β-Actin Monoclonal (Sigma Aldrich A1978). Incubation with HRP-conjugated secondary antibodies was performed for 1 hour prior to resolution with Immobilon Clasico HRP substrate (Millipore) on a BioRad Chemidoc imaging system. Quantification was performed using ImageJ. Secondary antibodies: Goat Anti-rabbit IgG, HRP-linked (Cell Signaling Technology 7074), Goat anti-mouse IgG (H+L) (Invitrogen 31430).

### RNA Sequencing

Total RNA was isolated from harvested ESCs using the Quick RNA Miniprep Kit, as per the manufacturer’s instructions (Zymo Research). RNA sample quality was confirmed by spectrophotometry (Nanodrop) to determine concentration and chemical purity (A260:A230 and A260:A280 ratios) before 1ug of each sample was sent for QC, library prep, and analysis at the Cornell Genomics Facility. Library preparation was performed using the NEBNext Ultra II [Directional] RNA kit with rRNA depletion (NEB). Libraries were sequenced to a minimum 20M raw reads on a NextSeq500 instrument (Illumina). For analysis, reads were trimmed for low quality and adaptor sequences. Reads were mapped to the mouse reference genome and gene expression analysis was performed using DESeq2.

### MTT Assay

Cell viability was analyzed using the CyQuant MTT Cell Viability Assay (ThermoFisher). Briefly, MTT was added to cells and incubated at 37°C for 4 hours before addition of SDS-HCl and incubation at 37°C for 18 hours. Following transfer of three technical replicates of resuspended dye solution to a 96-well plate, absorbance values (570nm) were obtained using a BioTek Synergy 2 plate reader. Technical replicates were averaged and normalized to vehicle-treated samples.

### Immunofluorescence Staining for Mitotic Abnormalities

Mouse ESCs were seeded in 8-well glass chamber slides (Millipore) 24 hours before treatment. Cells were treated for 24 hours prior to fixation with 4% formaldehyde for 10 minutes. Cells were then initially permeabilized with 0.25% Triton X-100 in phosphate-buffered saline (PBS-T) for 5 minutes, then fixed in ice-cold 100% methanol for a minimum of 30 minutes. Following rehydration with PBS, blocking was performed with 2% BSA and 5% goat serum in PBS (BS/GS) for 60 minutes at room temperature. Subsequently, incubation with mouse anti-phospho-Histone H2A.X (Ser139) clone JBW301 (Millipore Sigma 05-636) was performed for 60 minutes at 37°C. Cells were washed three times in PBS-T for 5 minutes before incubation with AF647-conjugated goat anti-mouse secondary antibody (Invitrogen A-21235) for 60 minutes at 37°C. Cells were again washed three times in PBS-T for 5 minutes before mounting with Prolong Gold Antifade Mounting reagent with DAPI.

Images were taken using a Zeiss LSM710. For each condition in each biological replicate, 20 pairs of cells were imaged and subsequently analyzed for mitotic abnormalities.

## Data availability statement

RNA sequencing data is available on NCBI GEO (Accession# GSE297363)

## Acknowledgements

This work was supported by grants R01HD082568 and P50HD096723 to JCS, and a training award from Cornell’s Center for Vertebrate Genomics to RCJ. We thank J. Grenier and the BRC Genomics Facility (RRID:SCR_021727) at the Cornell Institute of Biotechnology for sequencing experiments. We also thank Cornell’s Imaging Facility (RRID:SCR_021741) for assistance with imaging. Flow cytometry was performed at Cornell’s Flow Cytometry Facility (RRID:SCR_021740).

## Notes

### Competing Interest Statement

The authors have declared no competing interest.

