## Supplementary figures for "Embryonic Stem Cell-Specific Responses to DNA Replication Stress"

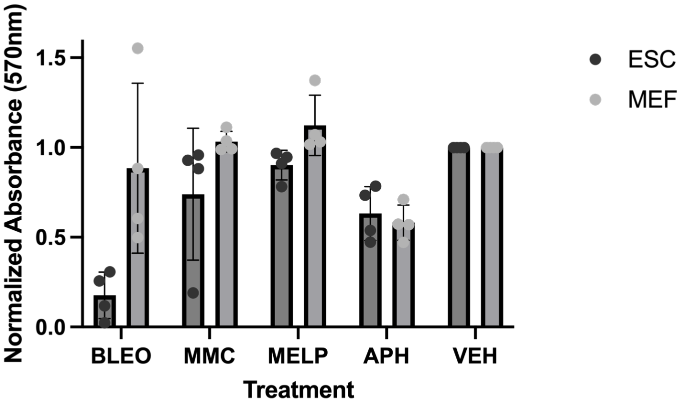


**Figure S1. Viability analysis via MTT of mESCs and MEFs treated with the indicated drugs for 24 hours.** Veh, vehicle; Bleo, bleomycin; MMC, mitomycin C; MELP, melphalan; APH, aphidicolin. Error bars show SD

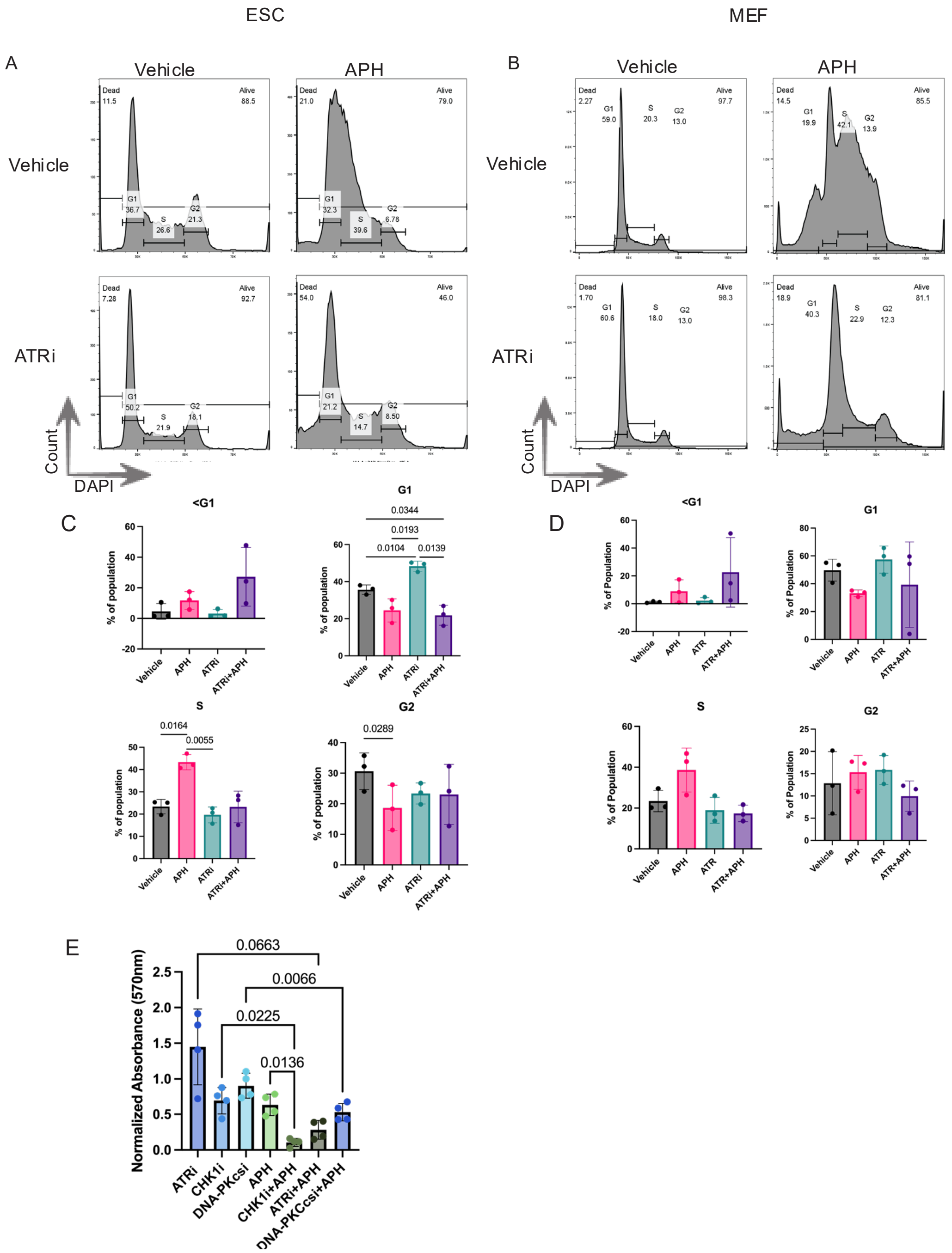


**Figure S2. Survival of ESCs under acute replication stress is ATR- and DNA-PKcs-dependent.** (A) Flow cytometry analysis of mESCs or MEFs (B) treated with the indicated drugs for 24 hours (C) Quantification of A (D) Quantification of B (E) Viability analysis via MTT of mESCs treated with the indicated drugs for 24 hours. All error bars show SD.

A

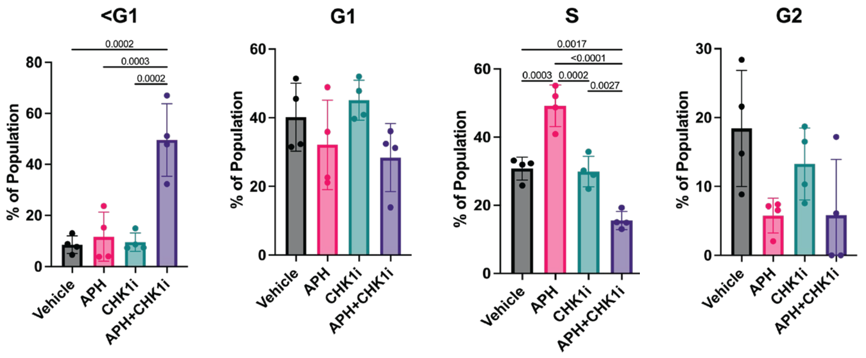


B

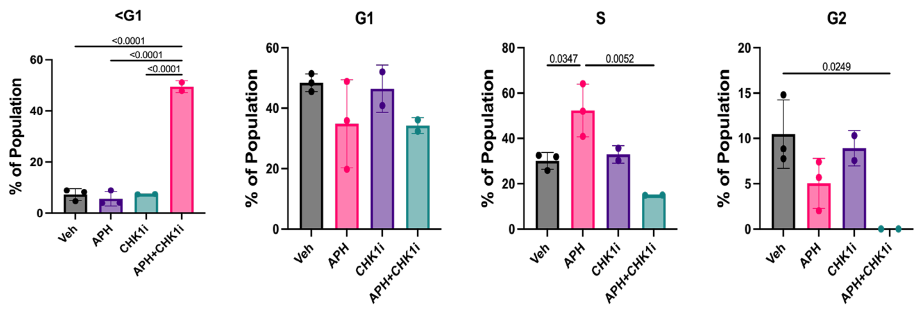


**Figure S3. Survival of cells under acute replication stress is CHK1-dependent** (A) Quantification of Figure 2A (ESCs). Error Bars show SD. (B) Quantification of Figure 2B (MEFs). Error Bars show SD.

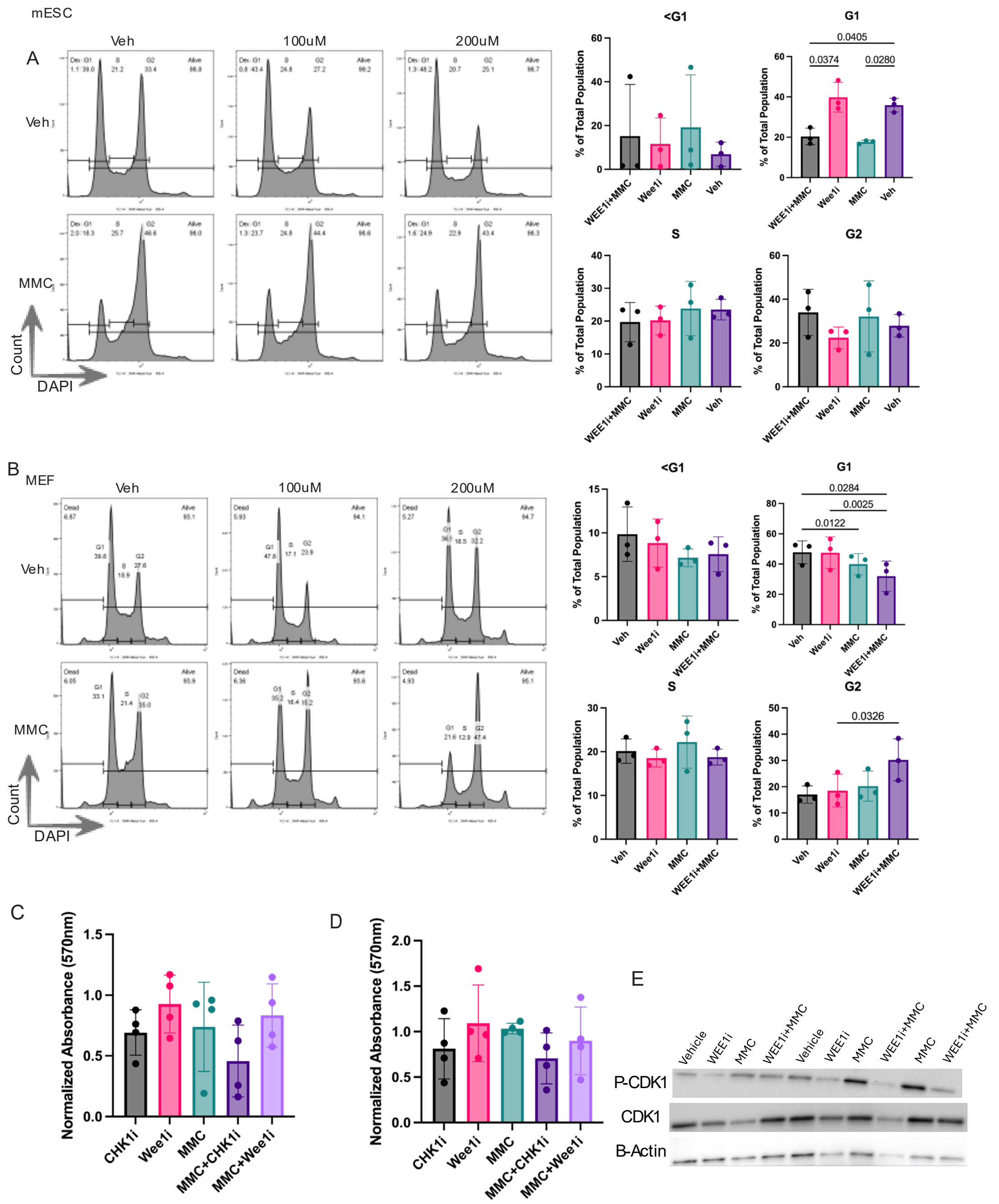


**Figure S4. Loss of WEE1 signaling induces different responses in mESCs and MEFs.** (A) Flow cytometry analysis of mESCs and MEFs (B) treated with Wee1i at the indicated concentrations and MMC for 24 hours. (C) Viability analysis via MTT assay of mESCs and MEFs (D) treated with the indicated drugs. (E) Western blot for p-CDK1 (Thr14, Tyr15) and CDK1 from MEFs treated with the indicated drugs for 24 hours. Multiple replicates shown. All error bars show SD

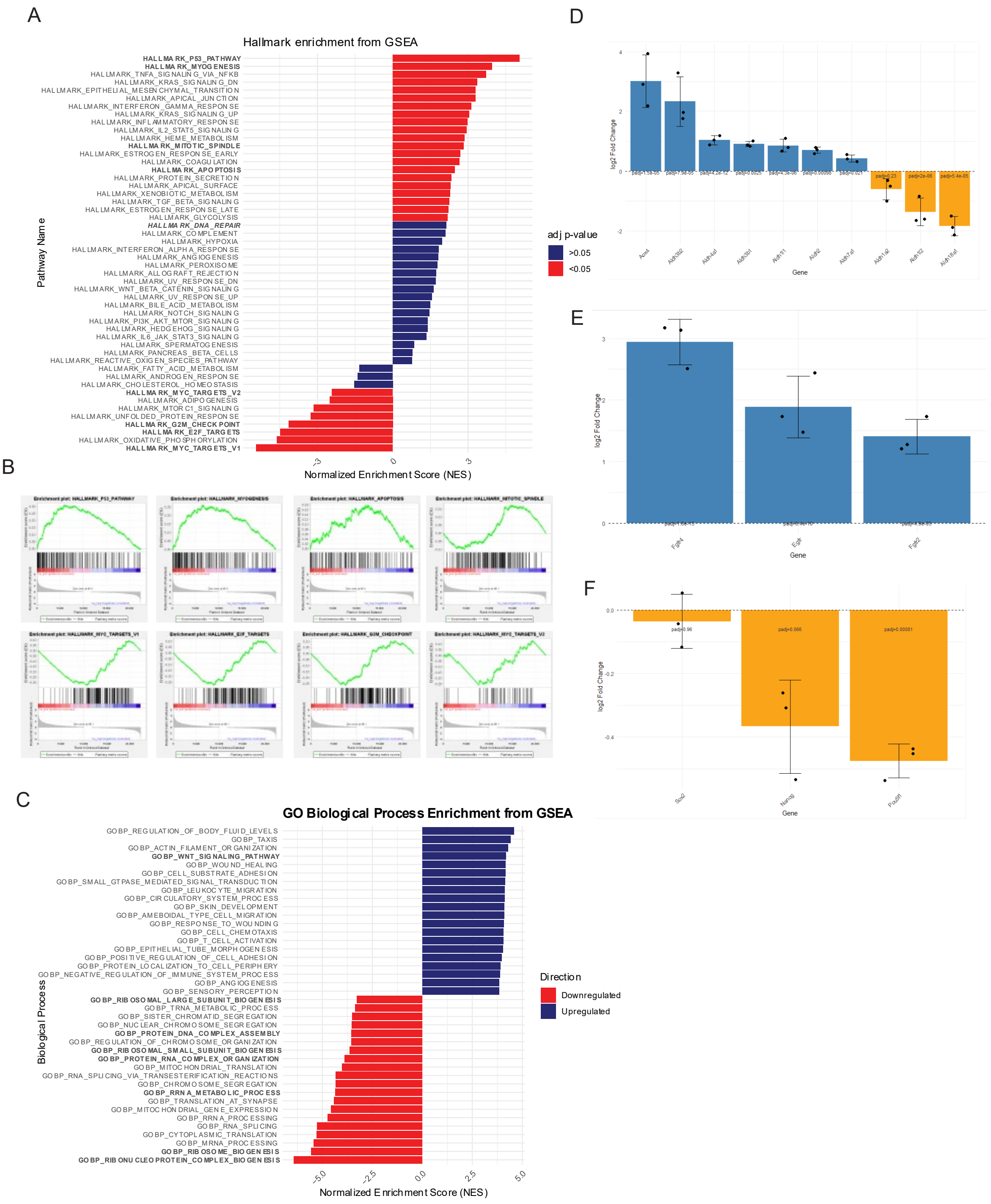


**Figure S5. MMC-induced transcriptional changes in mESCs.** (A) Hallmark enrichment plot (MMC-treated vs Vehicle) of all hallmarks from RNA sequencing of mESCs treated with MMC for 24 hours. (B) Representative enrichment plots. (C) GO Biological Process Enrichment of the top and bottom 20 most enriched processes from RNA sequencing of mESCs treated with MMC for 24 hours. (D) $Log_2$-Fold change of Aldehyde Dehydrogenase genes. (E) $Log_2$-Fold change of FGFR4, EGFR, and FGFR2. (F) $Log_2$-Fold change of Pluripotency genes. All error bars show SD.

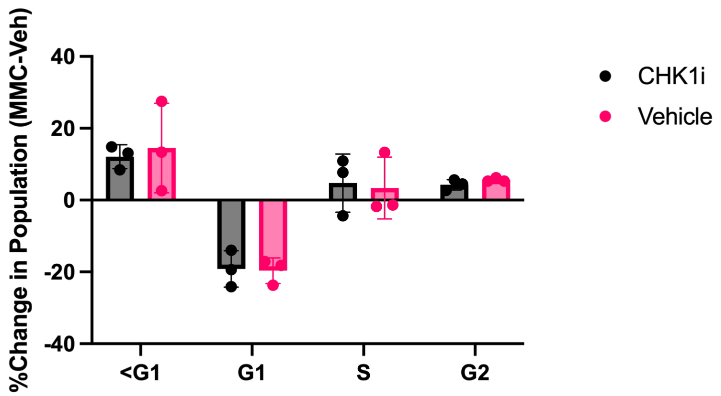


**Figure S6. CHK1i induces minimal changes in the response of mESCs to MMC** (A) Quantification of Figure 3D with MMC-treated samples normalized to vehicle. Demonstrates minimal CHK1i-dependent sensitization to MMC. Error bars show SD.

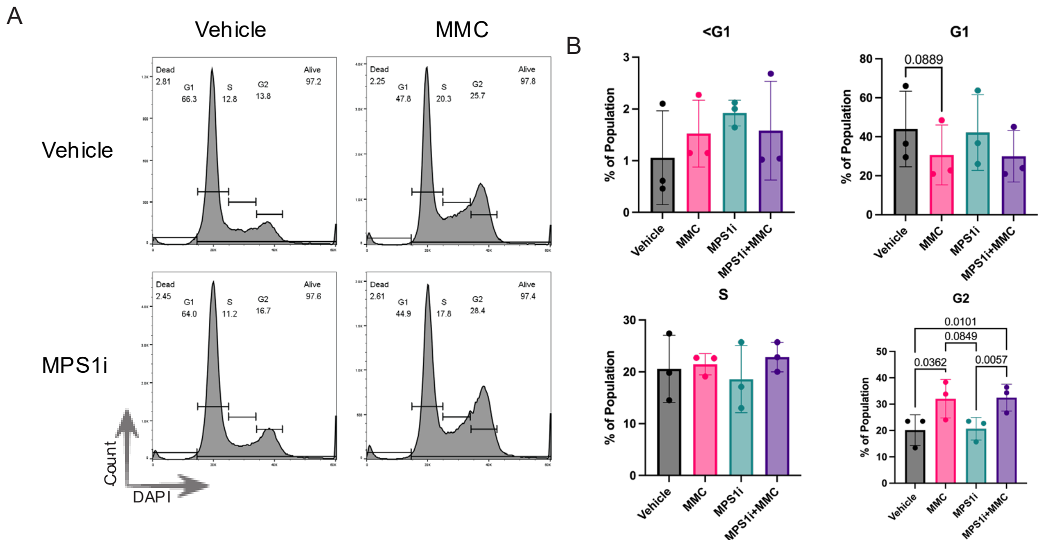


**Figure S7. MPS1i exerts no change in the response of MEFs to MMC.** (A) DNA content analysis from MEFs treated with the indicated drugs for 24 hours. (B) Quantification of A.

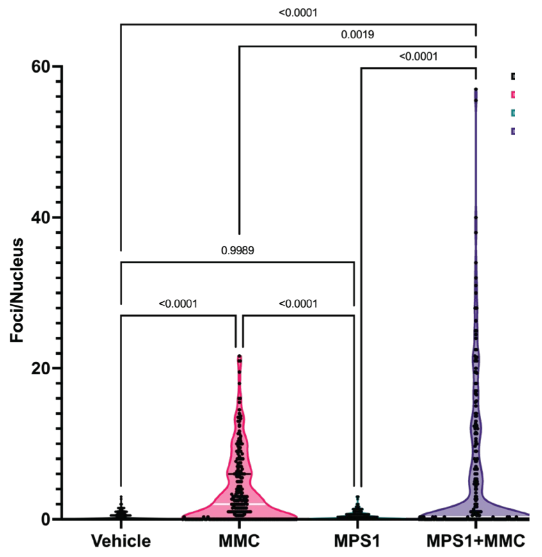


**Figure S8. Loss of the Spindle Assembly Checkpoint exacerbates MMC-induced genomic instability.** Quantification of yH2A.X foci per nuclear slice. Z=3, n>150, N=3.

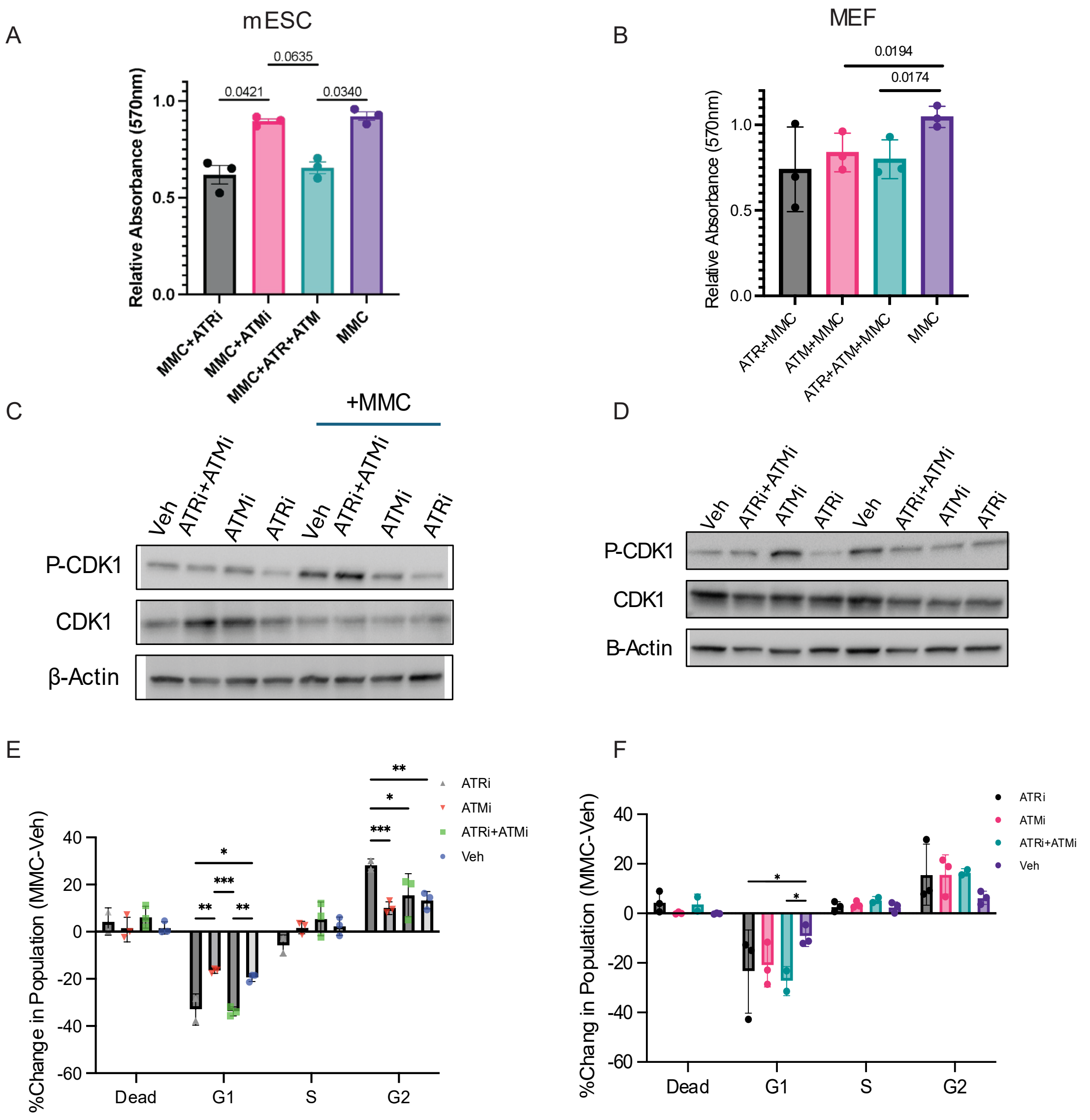


**Figure S9. Differential responses of ESCs and MEFs to the inhibition of ATR and ATM.** (A) MTT analysis from figure 5A with MMC-treated samples normalized to their corresponding vehicle-treated samples (B) MTT analysis from figure 5B with MMC-treated samples normalized to their corresponding vehicle-treated samples (C) Western blot analysis of mESCs treated with the indicated drugs. (D) Western blot analysis of MEFs treated with the indicated drugs. (E) Quantification of Figure 5D with MMC-treated samples normalized to their corresponding vehicle-treated samples (F) Quantification of Figure 5F with MMC-treated samples normalized to their corresponding vehicle-treated samples. All error bars show SD.
